# Predicting the Purity of Multispecific Antibodies From Sequence Using Machine Learning: Methods and Applications

**DOI:** 10.1101/2023.12.05.570217

**Authors:** Aviva Mazurek, Athena Davis, Stephen R. Comeau, Kenny Tsang, Javier Rivera, Zhong-Fu Huang, John Holt, Sandeep Kumar, Srinath Kasturirangan

## Abstract

Multispecific antibodies are prominent therapeutic agents, but many molecular formats and drug candidates that show promise during molecular discovery stages cannot be scaled up and developed into drugs due to inadequate developability. During the discovery stages, the selection of molecule format(s), molecule design, purity, and initial physiochemical stability testing criteria largely rely on scientists’ experience. Machine learning, however, can identify hidden trends in large datasets, aiding in the selection of drug candidates with improved developability. In this study, we present a machine learning approach to predict antibody purity, measured by the percentage of monomer after protein A purification. Using the amino acid sequences of variable regions, molecular formats, germlines and germline pairings, and calculated physiochemical properties as inputs, machine learning models were trained to predict the percentage of monomer for a given multispecific antibody (Figure 1). The dataset employed in this study consists of ∼500 multi-specific antibodies generated during BI’s internal drug discovery programs. Our results indicate that machine learning, when applied to sequence, germline, and format data, can effectively predict antibody percentage of monomer. Incorporating this approach into high-throughput multispecific antibody screening processes can save time and resources by reducing the need to test a large subset of potentially unstable antibodies. While this study focused on percentage of monomer as a test case, similar approaches can be employed to predict other antibody properties, such as melting temperature (Tm), hydrophobicity (aHIC), and solution stability properties (AC-SINS).

## 1. Introduction

Bispecific and multispecific antibodies are increasingly becoming commonplace as therapeutic modalities, particularly in Oncology^1^. Despite their widespread applications, multispecific antibodies suffer from developability and manufacturability issues. Several molecules that have shown potential in preclinical studies could not proceed to the clinic as they could not be scaled up, manufactured, or developed as a drug for patients^2^.

Typically, for a bispecific antibody to proceed to the clinic, it must have titers and monomer content that can ensure yields and quality^3^. Most multispecific antibodies that do not meet manufacturability threshold suffer from low yields and preclude their development into viable drugs^4^. Researchers often rely on empirical high throughput screening approaches to identify antibodies with the best developability profile. While it has been possible to produce and screen a relatively large panel of monoclonal antibodies, mixing and matching antibodies corresponding to 2 target binding arms to generate a bispecific is often limited by capacity issues of how many constructs can be cloned, expressed, purified, and tested^5^. Furthermore, while molecules may be very well behaved and show good developability properties as a monoclonal antibody (IgG), when these same sequences are paired together in a multispecific, it has been observed that the developability parameters of the monoclonal antibody do not always translate to the multispecific. Due to these limitations, researchers have largely relied on trial-and-error to empirically design and construct bispecific antibodies from parent IgG molecules which are then evaluated individually for their developability and function. These restrictions significantly limit multispecific antibody discovery and turnaround time to identify a viable lead candidate. These issues makes the design and testing of multispecific antibodies even more complicated^5^.

Machine learning has recently been incorporated into research and development, and specifically over the last few years in the realm of antibodies. Chen et al used machine learning to predict antibody developability from sequence. In order to do this, the authors looked at the antibody’s hydrophobic and electrostatic interactions as inferred from its three-dimensional structure and tried to correlate this to the antibody’s developability^6^. Liu et al used machine learning to generate models of antibody affinity that can create novel sequences with increased target specificity^7^. Li et all used machine learning to address the challenge of identifying features of antibodies that may influence antibody function^8^.

In this work, we present a machine learning approach to predict multispecific antibody purity, defined in terms of its percent monomer after Protein A purification, from the antibody sequence, format, physiochemical properties, and germline combinations. We designed this project as a simple classification problem and created a cutoff for a passing and failing percent monomer at 90%. Anything equal to or above 90% monomer is considered a passing antibody (“normal risk”), and anything below 90% monomer is considered a failing antibody (“high risk”). This process is illustrated in Figure 1.

We first trained our model on a training set consisting of percent monomer data for ∼500 multispecific antibodies. In addition to regular train-test split testing on this data, we further evaluated our model on three real world test sets and our models achieved 81-86% accuracy.

While our current analysis may be confounded by limited size of the data set and data bias, this study suggests that machine learning applied to sequence, germline, and format has the potential to be predictive of percent monomer and other biophysical properties such as melting temperature (Tm), hydrophobicity (aHIC), and solution stability properties.

## 2. Methods

The methods and calculations described in the following sections are demonstrated in the provided GitHub resource (amazurek1/ppma (github.com)).

This project was formulated as a supervised machine learning problem, utilizing sequence, physiochemical properties derived from homology modeling^9^, germline information, and antibody format as input features. The output is binary, represented by either “0” or “1”, where “1” signifies that the antibody has a monomer percentage equal to or greater than 90%, and “0” indicates a monomer percentage below 90%. We mined our internal production dataset to collect information on multispecific antibodies that have been produced at BI. The dataset comprises 212 antibodies with a monomer percentage equal to or greater than 90% (“normal risk”) and 271 antibodies with a monomer percentage below 90% (high risk). The information that was used as input to the model to train it to predict the percent monomer are described below.

### 2.1 Antibody Format

Multispecific antibodies can come in many different formats. At BI, our multispecific largely fall into 2 format categories – “Doppelmabs” and “Zweimabs”^10^. For this effort we focused on 2 Zweimab formats – Format A and Format B, and 1 Doppelmab format – Format C (Figure 1).

The Zweimab formats A and B represents an asymmetric format comprising 2 pairs of heavy and light chains which are brought together by knob-in-hole (KIH) mutations in the Fc. Format A has single chain Fab (scFab) domains on the knob and hole that comprise of the light chain region attached to the heavy chain region by a 38 amino acid flexible linker. Format B has a scFab on the knob and a single chain Fv (scFv) on the hole. They are typically monovalent for each target binder^11^.

The Doppelmab format, Format C, comprises of 2 Fabs that bind to the first target and 2 scFv that bind the second target. The scFv is appended to the C-terminus of the heavy chain. It is a symmetric format and is typically bivalent for each target binder^12^.

Each of these antibody formats were transformed into features, and the value for each of these features represents whether that antibody is embodied in that format (value of “1” or “0” depending on the formats presence or absence). Also known as “dummy variables”^13^.

In this dataset, there were 257 antibodies with Zweimab formats and 226 antibodies with Dopplemab formats (Figure 3a). The Zweimab formats were further categorized into Format A and Format B (Figure 3b).

**Figure 1.**
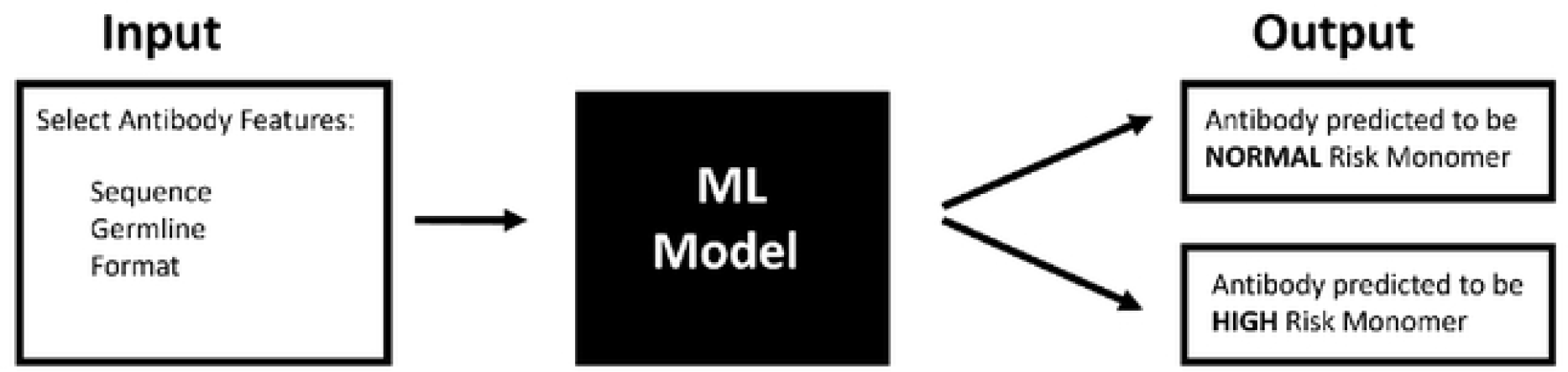
Overview of ML model for predicting multispecific antibody purity from sequence, germline and format information.

**Figure 2.**
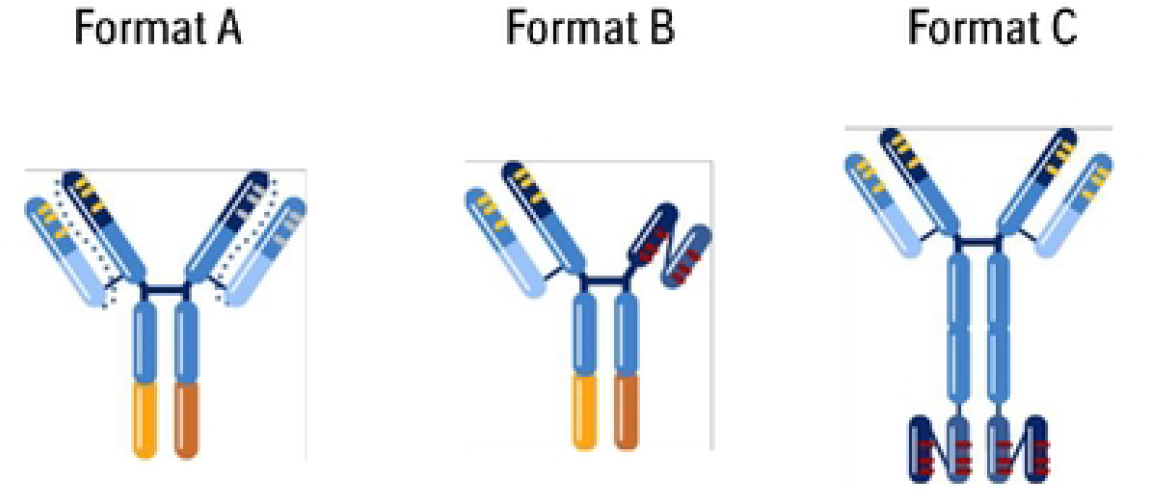
Description of formats and germlines used for ML training and test sets: Formats A and B - Zweimab formats; Format C - Doppelmab format, add information about germlines and germline pairs.

**Figure 3.**
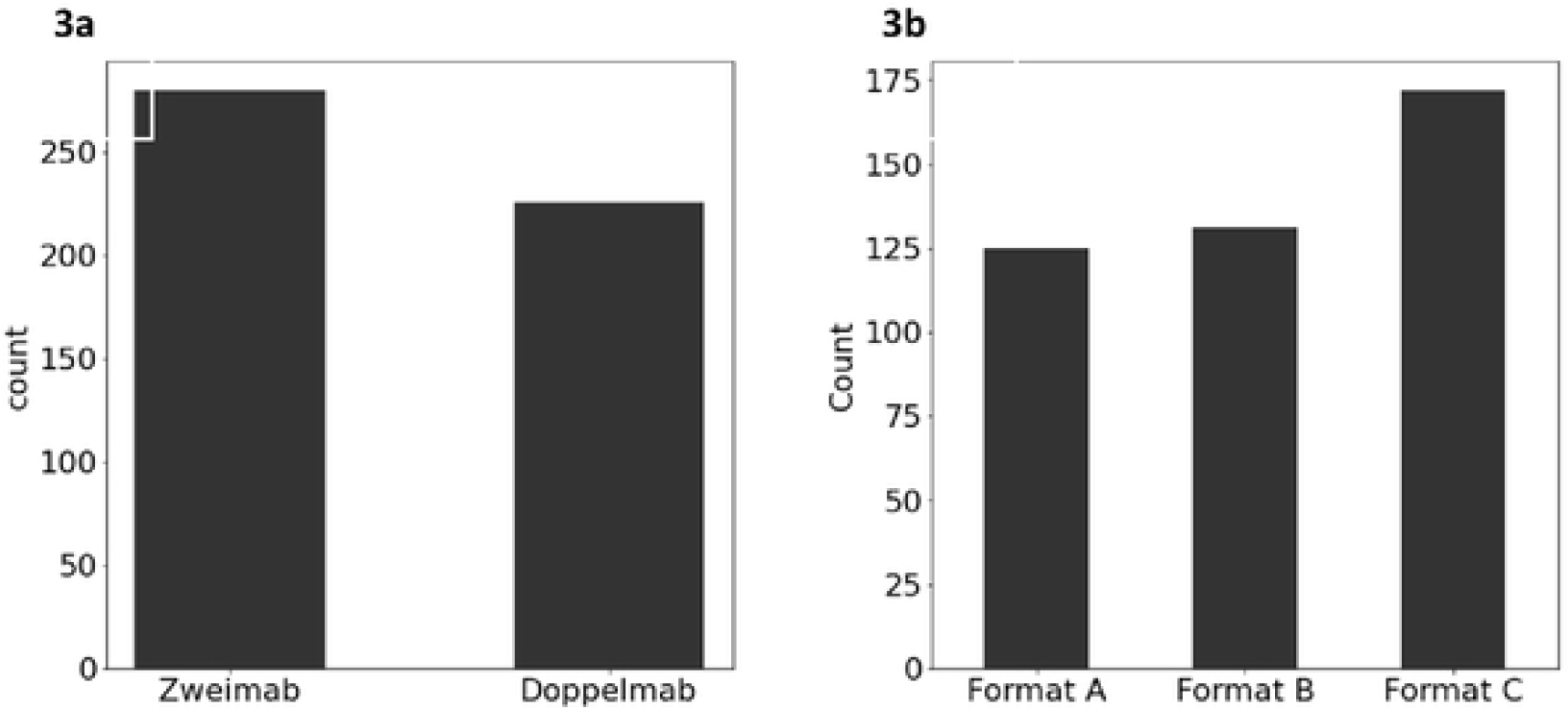
Distribution of multispecific formats used in the training data set. **3a)** Bar graph illustrating the composition of the training dataset consisting of 257 Zweimab and 226 Doppelmab antibodies. **3b)** Distribution of Zweimabs formats A, Band Doppelmab format C within the training dataset, as described in Figure 1.

### 2.2 Sequence

In order to extract features representing the amino acid distribution across the various chains of the antibody, we divided the antibody sequence into multiple features. First, the overall amino acid percentage of the 20 amino acids were computed and transformed into features for each antibody entry in the dataset, defined as “a” in Table 1. Twenty features were added into the dataset representing the 20 amino acids. For each antibody in the dataset, a value for each of these 20 features was calculated as a percentage between 0 and 100. This represents the frequency that each amino acid is present in the entire antibody.

**Table 1.** Dataset abbreviations and their corresponding names, serving as a quick reference guide for interpreting the heatmap in Figure 5. The abbreviations include “v” for amino acid percentages in individual framework and CDR components (using Kabat definitions), “a” for overall amino ac d percentages in the entire antibody dataset, “s” for the germlines dataset, “p” for the germline pairs dataset, “f” for the format of the antibody dataset, and “b” refers to physiochemical descriptors defined in supplementary Table 1.

| Dataset Abbreviation | Dataset |
| --- | --- |
| v | Amino acid percentage in individual frameworks and CDR components (using kabat definition) |
| b | Physiochemical Properties |
| a | Overall amino acid percentages in antibody |
| s | Individual Germlines |
| p | Germline Pairs |
| f | Format |

Additionally, each variable heavy and light chain was annotated according to its individual framework and CDR components using the Kabat nomenclature^14^. The antibody sequence was split up based on the individual framework and CDR components and was designated as “v” in Table 1. The amino acid percentages were computed for each of the framework and CDR regions across all of the variable domains (2xHeavy Chains, 2xLight Chains) of the multispecific antibody which resulted in 560 features per antibody (4 sequences * 7 regions (Framework 1-4, CDR1-3) * 20 amino acids).

### 2.3 Germlines

Antibody molecules are encoded by several recombined germline gene segments that provide antibodies with conformational flexibility and allows them to bind to many different antigens much like how one glove can fit the shape of many hands^15^.^16^ Each variable region in a given antibody sequence is wrapped in a generalized “germline” category^16^. The germlines corresponding to the variable heavy and light chain sequences were derived from IgBlast and incorporated into our model by converting them into “dummy variables”^13^. These values are either a “0”, “1”, or “2” in our dataset because a given germline is either not in the antibody (“0”), is present once in the antibody (“1”), or is present twice in the antibody(“2”). The overall value of the feature in the data frame for each multispecific antibody reflects how many times that germline was present in that antibody (referred to as “s” in Table 1).

For each multispecific antibody, the “Germline Pair” information was also computed. The “Germline Pair” corresponds to the exact pairing of the heavy and light chain germlines in the antibody. The germline pairing information was converted into dummy variables where a given multispecific antibody was either assigned a “0” (germline pair not present) or a “1”(germline pair present) in the feature set, referred to as “p” in Table 1.

For our training data set, the germline pairings and the frequency of occurrence of these germline pairs are shown in Figure 4a.^11^

**Figure 4.**
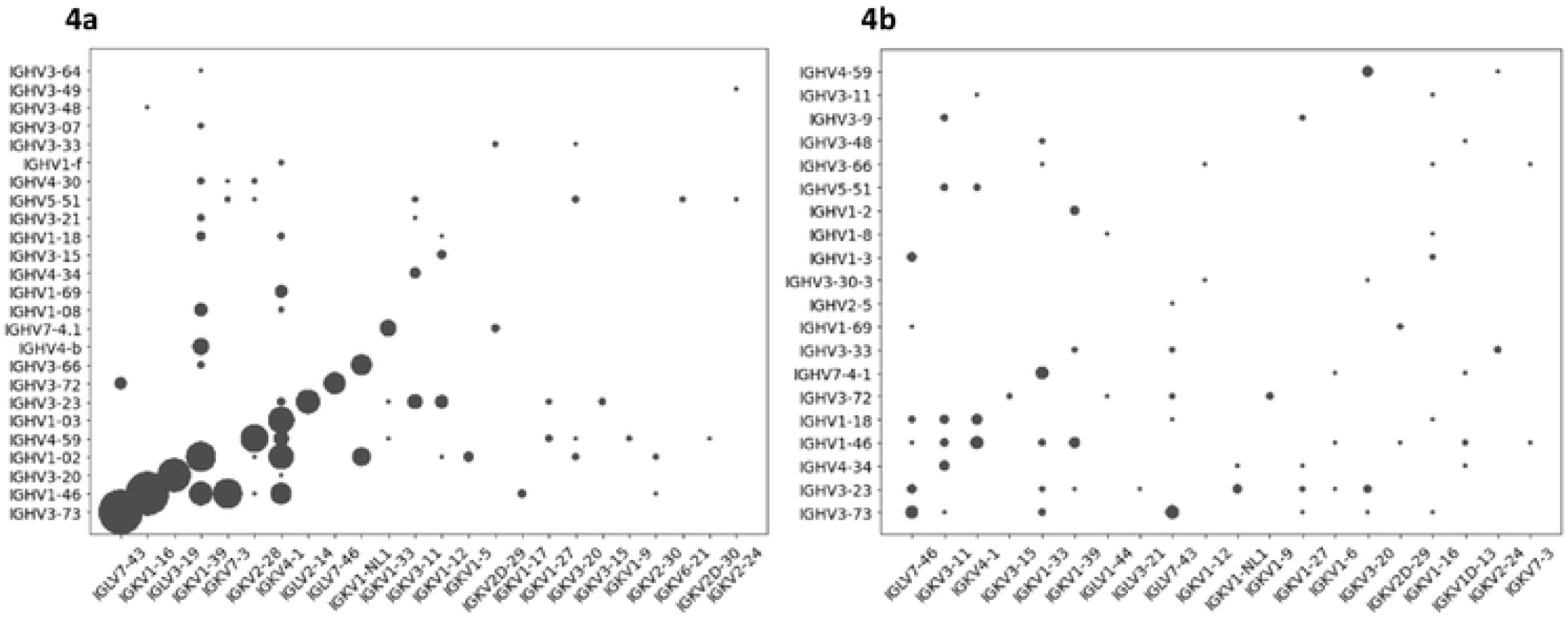
Scatterplots showing diversity of germline pairings in the training and test datasets. X-axis represents the light chains and y axis represents the heavy chains. **4a)** Represents diversity of germline pairings in the original training dataset of ∼500 antibodies **4b)** Diversity of germline pairings in test set 3.

## 3. Experimental Setup for Machine learning

The data corresponding to “v”, “a”, “s” and “f” described above were categorized into 6 distinct datasets: 1) the percentage of amino acids in the entire antibody (“v”), 2) the percentage of amino acids in individual framework and CDR components, as per the Kabat definition (“a”), 3) germlines (“s”) 4) germline pairs (“p”), 5) format (“f”). We also included an additional parameter 6) “physicochemical properties – “b” which corresponds to the physiochemical descriptors as described in Ahmed et al 2021^9^. These descriptors are listed in supplementary Table 1.

These datasets were amalgamated into all feasible combinations for 1,2,3,4,5, and 6 combinatorics, yielding 63 combinatoric datasets for testing (6 datasets for 1 combinatoric, 15 for 2 combinatoric, 20 for 3 combinatoric, 15 for 4 combinatoric, 6 for 5 combinatoric, and 1 for 6 combinatoric). These 63 dataset combinations are illustrated in the rows of **Figure 5**.

**Figure 5.**
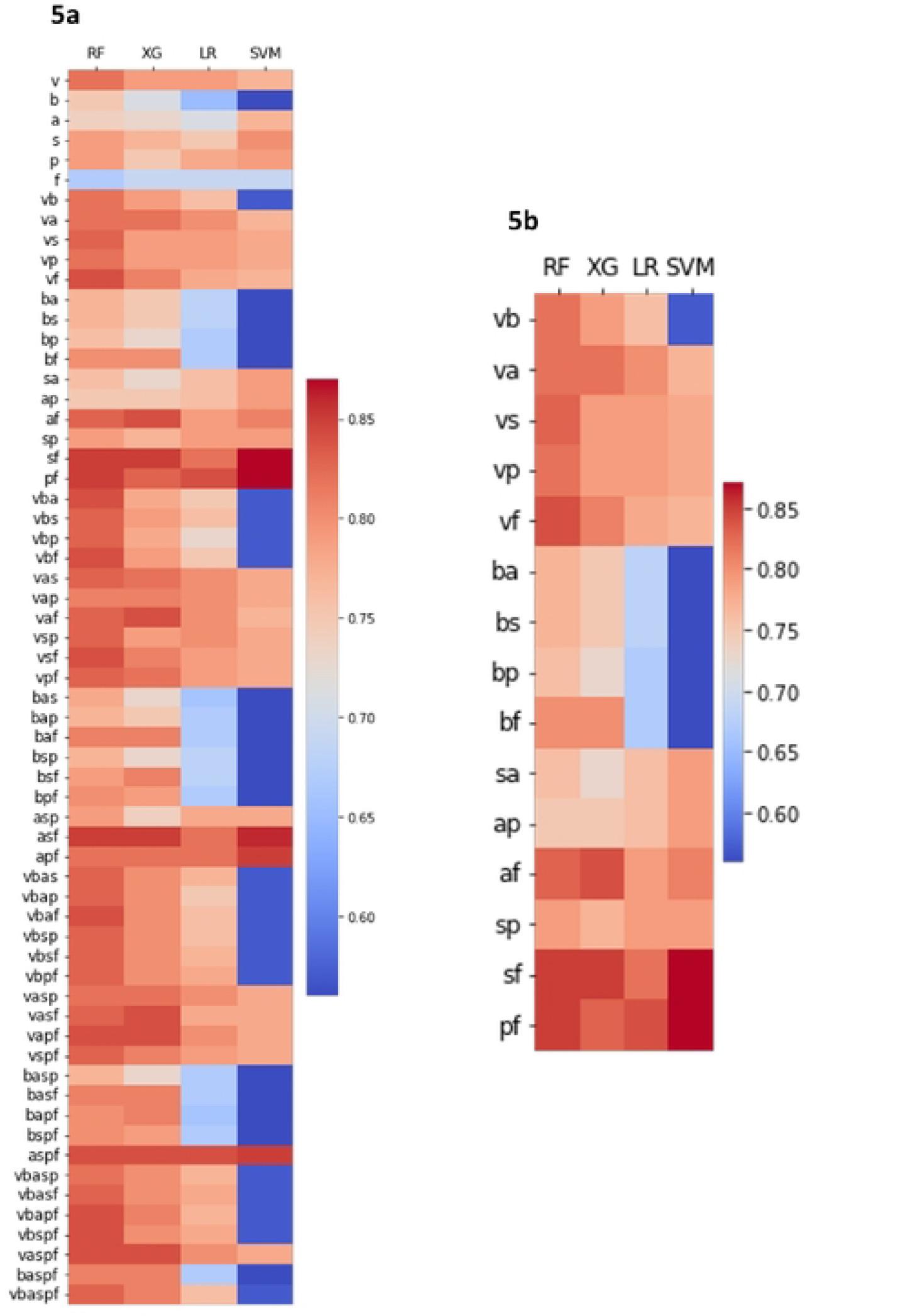
Performance of different machine learning models across different feature set combinations. Color intensity from dark red to dark blue indicates highest to lowest accuracy **5a)** Heatmap illustrating the cross validation (CV) accuracy scores for each feature, feature combination (ranging from 1 to 6 combinator c), and four ML models (RandomForest, XGBoost, Logistic Regression, and Support Vector Machine) across the six datasets. Each row represents a unique combination of datasets, and each column represents an ML model. Refer to Table 2 for dataset abbreviations and their corresponding names. **5b**) Excerpt from 5a focused on 2 combinatorics shows 1. RF (column 1) outperforming other ML models 2. Importance of format feature (f) for enhanced model accuracy.

To train and assess these 63 dataset combinations, we initially executed an 80/20 stratified train-test split on each combination. Subsequently, each combination was trained using scikit-learn with RandomForest, XGBoost, Logistic Regression, and Support Vector Machine models. The performance of each model was evaluated using a 10-fold cross-validation accuracy for comparison. This experiment facilitated the comparison of the performance of various models and dataset combinations, enabling us to identify the most predictive models and dataset combinations (Figure 5).^17,18,19^

**Table 2.** summary statistics for comparison of ML models for %monomer experiments. First columns lists the given model, and the second column is how often the model outperforms all other models in the experiments. Percentages were rounded to the nearest ones place.

| Model | Outperforms All Other Models (%) |
| --- | --- |
| Random Forest | 63% |
| XGBoost | 18% |
| Logistic Regression | 0% |
| Support Vector Machine | 18% |

In addition, subsequent to the production phase, we subjected our final model to three real-world test sets. These test sets served to evaluate the performance of our model in a real-world context.

### 3.1 Pressure test datasets

In order to pressure test the ML algorithm, we ran 3 real world test sets. The first test set consisted of 15 multispecific antibodies, the second incorporated 75 multispecific antibodies, and the third encompassed 90 multispecific antibodies. For all 3 test sets, we used different combinations of Formats A, B and C and produced the molecules in the lab and tested them for percent Monomer after Protein A purification.

The germlines used in the first two test sets were more similar to our training data, providing a more direct comparison of the model’s predictive capabilities.

To further pressure test the ML algorithm, we combined the variable heavy and light chain sequences of 37 clinically validated bispecific antibodies to generate over 4,000 unique combinations in Format X and Y. We then applied the ML algorithm to analyze these antibodies and based on the predictions of high and low risks, selected a diverse set of 90 antibodies for laboratory testing to validate the results. This test set included 122 new germline combinations and 58 combinations that the model had previously encountered during training. Test set 3 represents 45 antibodies where both germline pairings that were not present in the training set, 77 antibodies that had at least one germline pairing that was not present in the training set, and 13 antibodies where both germline pairings were present in the training set.

This diversity enabled us to further investigate the model’s performance across a broader spectrum of germline pairings and effectively test it in varying scenarios. The germline diversity used in test set 3 is shown in Figure 4b.

## 4. Experimental Setup for Antibody Production and Characterization

### Multispecific antibody design

All multispecific antibodies in Formats A, B or C were constructed by recombinant DNA technology. Genes encoding variants of binders to Target 1 or Target 2 were obtained by gene synthesis of light chain and heavy chain variable domains (Genewiz). The human IgG1 Fc with effector function knock-out (LALA) mutations was used as Fc-region for all multispecific antibodies. The molecules in Formats A and B contained Genentech’s knob-in-hole mutations in the CH3 domain to promote heterodimerization^20^.

### Multispecific antibody expression and purification

All multispecific antibodies were expressed from transfected Chinese hamster ovary (CHO) cells and expressed and purified using Protein A chromatography as previously described^21^.

### Size exclusion chromatography (SEC)

In order to determine the % monomer of the multispecific antibodies, analytical size exclusion chromatography was carried out using the Agilent HPLC system as previously described^22^. 10&g of sample were injected onto two tandemly connected Acquity BEH200 columns (Waters, 1.7 &m, 4.6 X 150 mm) at a flow rate of 0.25 mL/min. The mobile phase contained 50 mM sodium phosphate, 200 mM arginine chloride, 0.05% sodium azide, pH 6.8. The elution profile was monitored at 280 nm and manually integrated to calculate the level of high molecular weight (HMW) and low molecular weight (LMW) species. Representative SEC chromatograms are shown in Figure 6.

**Figure 6.**
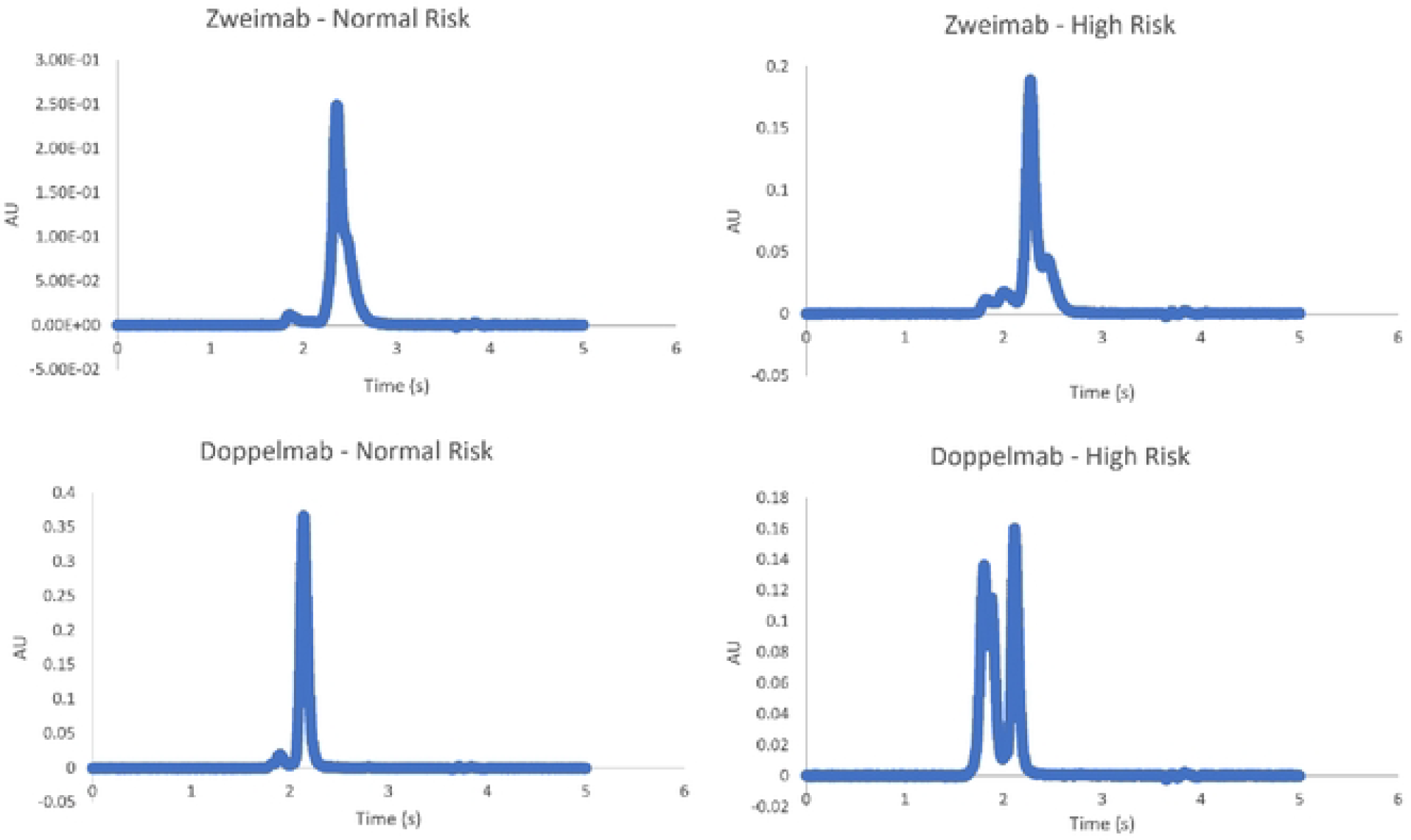
**Representative analytical size exclusion chromatography** results for Zweimab and Doppelmab showing molecules with normal and high risk behavior

## 5. Results and Discussion

The studies described herein have been designed to determine if machine learning could assist in identifying antibody combinations which would be manufacturable in the lab.

First, we tried to determine if the format or germline by themselves influenced the risk profile of the multispecific antibodies used in the learning data set.

Figure 7 displays representative examples of monomer % of multispecific antibodies across a pair of germlines and different formats. This data indicates that for a given pair of germlines and a given format (Doppelmab or Zweimab), there is a range of monomer %, indicating that the germline and format definitions alone does not seem to be predictive of the purity, or indicative of a trend for %monomer.

**Figure 7.**
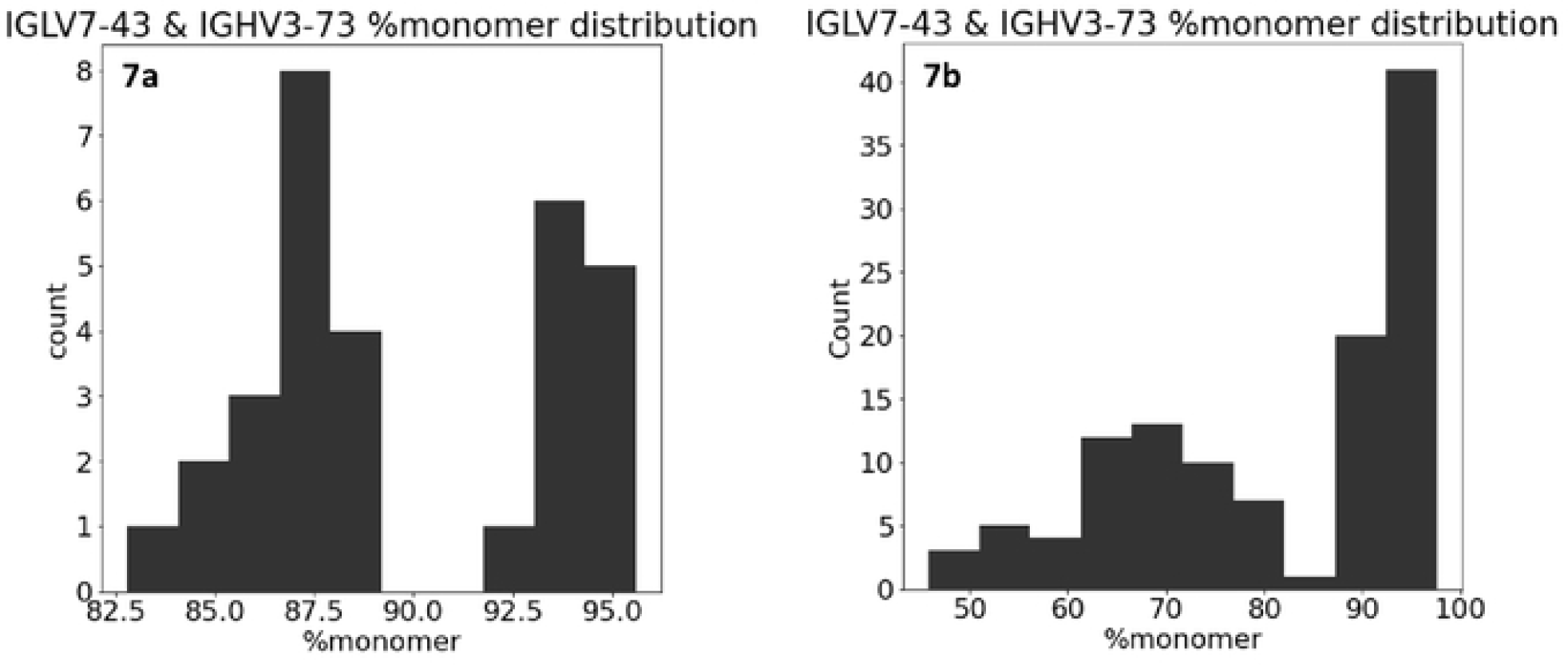
Bar graphs illustrating the distribution of %monomer and count for different formats and germline combinations. The X axis represents %monomer, and the Y axis represents count of each combination of format and germline pair in the dataset in both 6ba and 6bb. **7a)** Format: Doppelmab and germline combination IGLV7 43 & IGHV3-73; **7b)** Format: Zweimab and germline combination IGLV7 43 & IGHV3-73.

The monomer distribution does not also exhibit any clear trend when particular germline pairs are analyzed together for a given format. For e.g. IGKV4-1 and IGKV1-39 are among the more prevalent germlines in the dataset, and the data for Zweimab and Doppelmab molecules across these germlines cover a range of monomer % indicating the absence of a specific trend that is governed by the germlines used in the dataset (Figure 7).

Given the lack of a clear trend in the %monomer distribution when specific germline pairs and formats are combined, it was necessary to implement machine learning models that take into account multiple features. This approach helps identify trends that incorporate various aspects of the data, which may not be immediately apparent through human analysis.

To better understand the performance of individual features, feature combinations, and the predictive power of the various machine learning (ML) models, we generated a heatmap illustrating the cross-validation (CV) accuracy scores for each feature, feature combination (ranging from 1 to 6), using four ML models for each dataset (Figure 5). The ML models evaluated included RandomForest, XGBoost, Logistic Regression, and Support Vector Machine.

In **Figure 5**, each dataset is represented by an abbreviation: “v” for the amino acid percentages in each individual framework and CDR component (using the Kabat definitions), “a” for the overall amino acid percentages in the entire antibody dataset, “s” for the germlines dataset, “p” for the germline pairs dataset, “f” for the format of the antibody dataset, and “b” refers to physiochemical descriptors that have previously defined in Ahmed et al 2021^9^.

To further clarify these abbreviations, a table detailing the dataset names corresponding to each abbreviation is provided in **Table 2**. This table serves as a quick reference guide, aiding in the interpretation of the heatmap and the understanding of the performance of the different features and ML models across the six datasets.

When comparing the results of the individual datasets, the amino acid percentages in the individual frameworks and CDR components perform the best on their own in predicting the risk for precent monomer. While the Format of the antibody has very low predictive power on its own, when combined in a two combinatoric with any other dataset it improves the accuracy more than any other pair.

Figure 5b illustrates all two-way combinations of datasets and the four ML model results. It is particularly noticeable in the Random Forest and XGBoost models that any combination involving the Format dataset results in higher accuracy.

Considering the overall results of Figure 5 we decided to proceed with Random Forest for several reasons. Firstly, it demonstrated the best overall performance. Random Forest outperforms the other models by a large majority. A summary of how often each model outperforms all other models is summarized in Table 3.

Random Forest outperforms all other models in each dataset combination by 63%. More importantly, it exhibits the highest level of consistency across various combinations of datasets, which is crucial for our project as the input data and the number of features will vary depending on the specific project circumstances. This consistency, coupled with its superior performance, reinforces our decision to proceed with the Random Forest model.

RandomForest is a robust ML technique that utilizes bagging and feature randomness^23^. It handles outliers very well by binning them, maintains a low bias in its decision trees, and handles high dimensionality well. Our training dataset has 447 datapoints, and up to 700 features depending on which features are included, which means it contains high dimensional data. Not only does RandomForest handle this type of data well, since it works with subsets of data, but in many cases it can actually improve the model with this type of data^23^.

We focused our research on the RandomForest model and feature combinations that align more closely with our ultimate goal or use case. We trained and fine-tuned the hyperparameters of the RandomForest model on ten different dataset combinations. We then calculated the cross-validation accuracies and test accuracies for each of these combinations. The results of these computations are illustrated in Figure 8. The overall trend indicates that the inclusion of more features from our dataset leads to an increase in accuracy.

**Figure 8.**
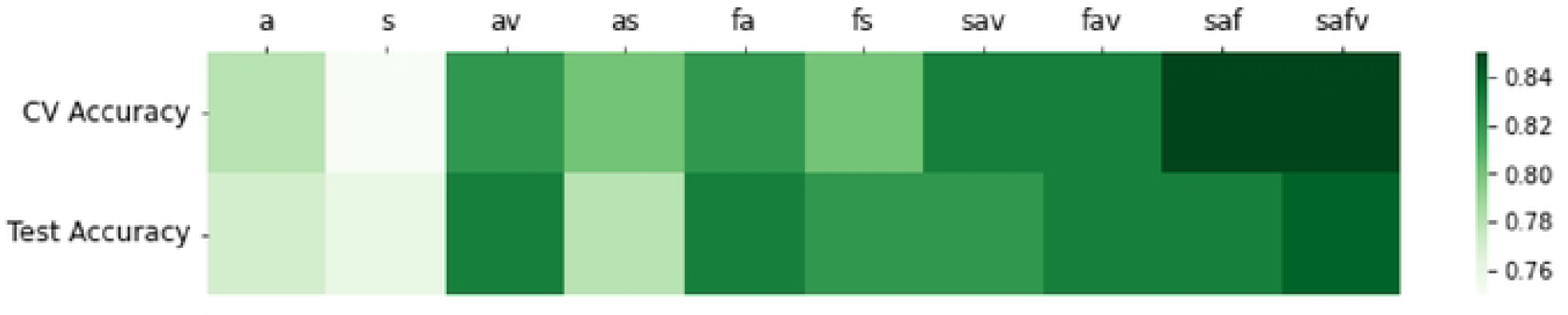
Heatmap displaying cross validation accuracies and test accuracies for predicting the percentage of monomer using the Random Forest model across ten distinct feature sets. Test accuracy refers to accuracy of RF model trained on a random train-test split predicting the outcome of that random test set.

The general trend in Figure 8 is that combining the germline, amino acid percent, format, and amino acid percentages of the Kabat regions data produces a more accurate model. This implies that distinct features within the model contribute unique information, thereby enhancing the model’s overall accuracy.

The model that incorporates all the features (safv) achieves an F1 score of approximately 83.3%. The confusion matrix for the test set from a random train-test split corresponding to this model is illustrated in Figure 9a. A confusion matrix is often used in the Machine Learning world to represent the performance of a classification model by displaying the number of true positive (TP), true negative (TN), false positive (FP), and false negative (FN) predictions in each quadrant. The top-left quadrant contains the TP values, the top-right quadrant contains the FP values, the bottom-left quadrant contains the FN values, and the bottom-right quadrant contains the TN values. This matrix provides a clear visualization of the model’s accuracy and potential areas for improvement.

**Figure 9.**
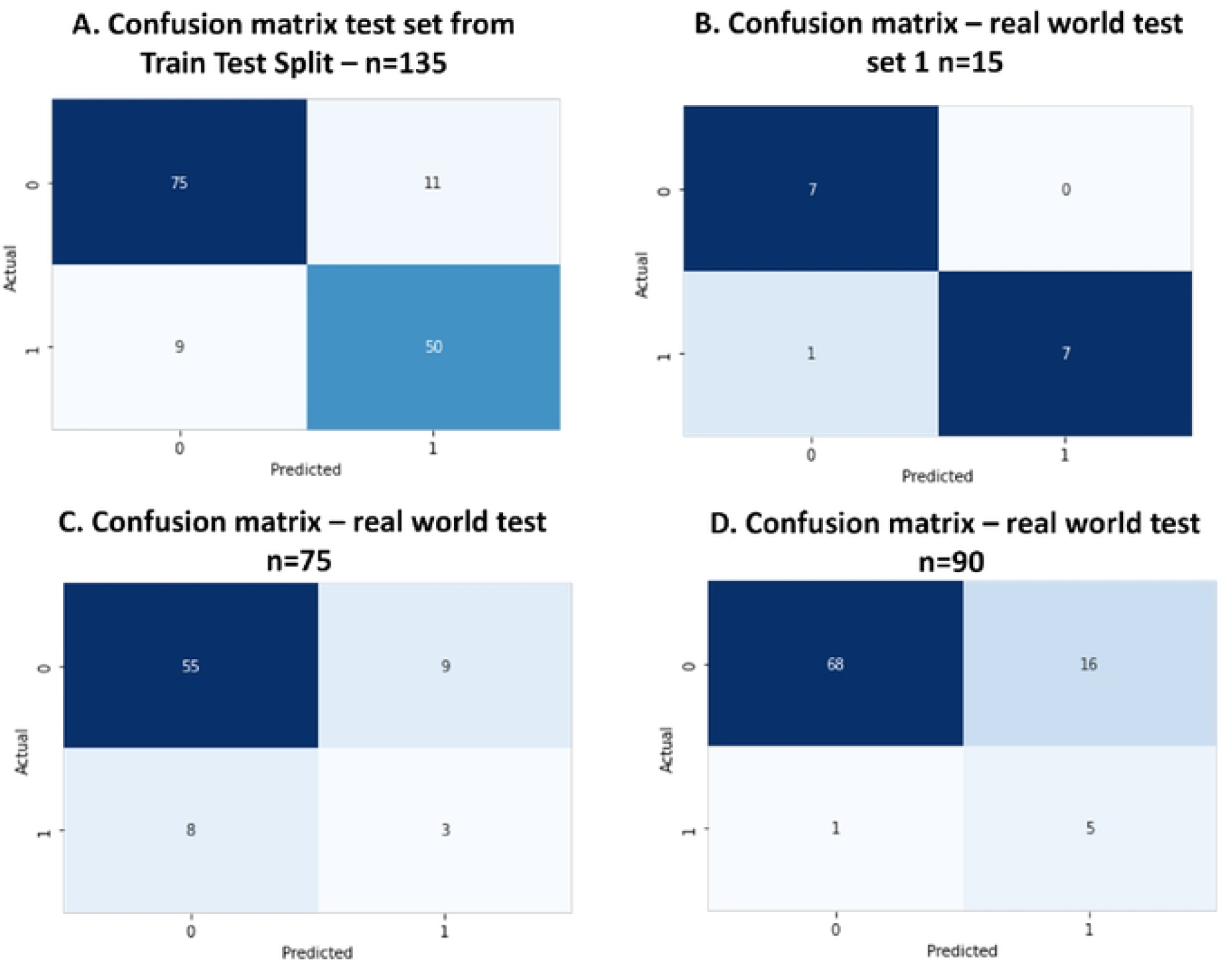
**9a)** Confusion Matrix for the test set for the random forest model that includes all features. The top left quadrant represents the number of times the model accurately predicted antibodies to be below 90% monomer. The top-right quadrant indicates the instances where the model incorrectly predicted antibodies to be equal to or above 90% monomer when they were actually below 90%. The bottom-left quadrant shows the cases where the model predicted antibodies to be below 90% monomer, but they were in fact equal to or above 90% monomer. Last y, the bottom-right quadrant represents the correct predictions of the model when antibodies were equal to or above90% monomer. **9b)** Confusion Matrix for first real world test set. **9c)** Confusion Matrix for second real world test set. **9d)** Confusion Matrix for third real world test set.

Finally, we applied the model to three real-world test sets. For all 3 real-world tests, we first predicted the outcome using the ML algorithm and then went ahead to produce and test these multispecific antibodies in the lab. We then compared the empirical data to the predictions, the results of which are represented by the three confusion matrices.

The first test was a small set of 15 molecules in Format B and yielded a confusion matrix with 7 true positives, 1 false positive, 0 false negatives, and 7 true negatives (Figure 9b).

The second test set was a larger set in Format B where one of the arms of the bispecific antibody was “fixed” to a sequence that had previously been used in the training set, and thus exposed to the ML algorithm, but the 2^nd^ arm comprised of new antibody sequences from very similar germlines. This set produced a confusion matrix with 55 true positives, 8 false positives, 9 false negatives, and 3 true negatives (Figure 9c).

Both test sets contained sequences that were similar to those in the training sets and had highly represented germlines in the training dataset. Therefore, the excellent performance of the model on these test sets was expected.

The third test set was more challenging. We designed 90 multispecific antibodies comprising of sequences that had not been used in the training set, and thus comprised of more diverse germlines than those in the previous test sets. Test set 3 represents 45 antibodies where both germline pairings were not present in the training set, 77 antibodies that had at least one germline pairing that was not present in the training set, and 13 antibodies where both germline pairings were present in the training set.

We ran them through the model before testing the percent monomer in the lab. The resulting confusion matrix had 68 true positives, 1 false positive, 16 false negatives, and 5 true negatives (Figure 9d). Of the six antibodies that were classified as normal risk, the model accurately predicted five of these as normal risk. Among these five antibodies, there were 10 germline combinations due to two branches. Eight of these combinations were entirely new, while two were combinations that the model had previously seen during training. Interestingly, the two antibodies with previously seen combinations also had a germline pairing that was new to the model.

The lower test accuracy for the third test set, can be attributed to the increased diversity of the sequences used in this dataset compared to the data used for training the model and in the first 2 test sets. This is a common challenge in machine learning and underscores the importance of having diverse and large training data sets.

Despite this, the model’s performance on the third test set demonstrates its potential to handle diverse antibody sequences. This model is still in its early stages. As we continue to collect more diverse data and continue retraining, it will progressively become more robust and reliable.

Overall, this approach has the potential to efficiently screen a vast number of multispecific antibody combinations, allowing us to eliminate candidates that would otherwise be unsuitable for testing in the lab. The ability to predict the suitability of an antibody for lab testing based on its sequence is a powerful tool that can save significant time and resources in the development of new therapeutics.

In conclusion, our model has shown promising results in its ability to accurately predict the suitability of antibodies for lab testing. As we continue to refine the model and expand our training data, we anticipate that its performance will continue to improve, making it an invaluable tool in the field of multispecific antibody development.

---

**Supplementary material Table 1.**

| **Code** | **Description** |
| --- | --- |
| Atom Clashes | Atom clash count (> 0.5kcal/mol) as from Homology Model |
| BSA_LC_HC | BSA between light and heavy chains |
| E | Potential Energy |
| Eint_VL_VH | Affinity interaction energy between VL and VH (same as affinity_VL_VH) |
| HI | Hydrophobic imbalance |
| Packing Score | Residue packing quality score as from Homology Model |
| Phi Psi Outliers | Backbone psi and phi angle outliers (Z > 4) |
| pro_asa_hph | Hydrophilic Surface Area |
| pro_asa_hyd | Hydrophobic Surface Area |
| pro_cdr_net_charge | Protein Net Charge on CDRs only |
| pro_dipole_moment | Protein Dipole Moment |
| pro_hyd_moment | Hydrophobicity Moment |
| pro_net_charge | Protein Net Charge |
| pro_patch_cdr_hyd | Area of hydrophobic protein patch(es) near CDRs |
| pro_patch_cdr_neg | Area of negative protein patch(es) near CDRs |
| pro_patch_cdr_pos | Area of positive protein patch(es) near CDRs |
| pro_patch_hyd | Area of hydrophobic protein patch(es) |
| pro_patch_neg | Area of negative protein patch(es) |
| pro_patch_pos | Area of positive protein patch(es) |
| pro_pl_3D | Structure-based pI Prediction |
| pro_pi_seq | Sequence-based pI Prediction |
| pro_zeta | Zeta Potential at Debye Length |
| Rotamer Outliers | Rotamers with strain > 5kcal/mol as from Homology Model |

## References

1. Brinkmann, U. & Kontermann, R. E. The making of bispecific antibodies. mAbs 9, 182–212 (2017).

2. Husain, B. & Ellerman, D. Expanding the Boundaries of Biotherapeutics with Bispecific Antibodies. BioDrugs 32, 441–464 (2018).

3. Underwood, D. J., Bettencourt, J. & Jawad, Z. The manufacturing considerations of bispecific antibodies. Expert Opin. Biol. Ther. 22, 1043–1065 (2022).

4. Bailly, M. et al. Predicting Antibody Developability Profiles Through Early Stage Discovery Screening. mAbs 12, 1743053 (2020).

5. Segaliny, A. I. et al. A high throughput bispecific antibody discovery pipeline. Commun. Biol. 6, 380 (2023).

6. Chen, X. et al. Predicting Antibody Developability from Sequence using Machine Learning. bioRxiv 2020.06.18.159798 (2020) doi:10.1101/2020.06.18.159798.

7. Liu, G. et al. Antibody Complementarity Determining Region Design Using High-Capacity Machine Learning. Bioinformatics 36, 2126–2133 (2019).

8. Li, X., Deventer, J. A. V. & Hassoun, S. ASAP-SML: An antibody sequence analysis pipeline using statistical testing and machine learning. PLoS Comput. Biol. 16, e1007779 (2020).

9. Ahmed, L. et al. Intrinsic physicochemical profile of marketed antibody-based biotherapeutics. Proc National Acad Sci 118, e2020577118 (2021).

10. Venkataramani, S. et al. Design and characterization of Zweimab and Doppelmab, high affinity dual antagonistic anti-TSLP/IL13 bispecific antibodies. Biochem. Biophys. Res. Commun. 504, 19–24 (2018).

11. Hipp, S. et al. A Bispecific DLL3/CD3 IgG-Like T-Cell Engaging Antibody Induces Antitumor Responses in Small Cell Lung Cancer. Clin. Cancer Res. 26, 5258–5268 (2020).

12. Coloma, M. J. & Morrison, S. L. Design and production of novel tetravalent bispecific antibodies. Nat. Biotechnol. 15, 159–163 (1997).

13. Wooldridge, J. M. (2016). Introductory Econometrics: A Modern Approach (6th ed.). Boston, MA: Cengage Learning.

14. Johnson, G. & Wu, T. T. Kabat Database and its applications: 30 years after the first variability plot. Nucleic Acids Res. 28, 214–218 (2000).

15. Willis, J. R., Briney, B. S., DeLuca, S. L., Crowe, J. E. & Meiler, J. Human Germline Antibody Gene Segments Encode Polyspecific Antibodies. PLoS Comput. Biol. 9, e1003045 (2013).

16. Jung, D. & Alt, F. W. Unraveling V(D)J Recombination Insights into Gene Regulation. Cell 116, 299–311 (2004).

17. Breiman, L. Random Forests. Mach. Learn. 45, 5–32 (2001).

18. Probst, P., Wright, M. N. & Boulesteix, A. Hyperparameters and tuning strategies for random forest. Wiley Interdiscip. Rev.: Data Min. Knowl. Discov. 9, (2019).

19. Zhu, N., Zhu, C., Zhou, L., Zhu, Y. & Zhang, X. Optimization of the Random Forest Hyperparameters for Power Industrial Control Systems Intrusion Detection Using an Improved Grid Search Algorithm. Appl. Sci. 12, 10456 (2022).

20. Spiess, C. et al. Bispecific antibodies with natural architecture produced by co-culture of bacteria expressing two distinct half-antibodies. Nat. Biotechnol. 31, 753–758 (2013).

21. Liu, Y. et al. An adapted consensus protein design strategy for identifying globally optimal biotherapeutics. mAbs 14, 2073632 (2022).

22. Liu, Y. et al. An adapted consensus protein design strategy for identifying globally optimal biotherapeutics. mAbs 14, 2073632 (2022).

23. Bubeck, S. & Sellke, M. A Universal Law of Robustness via Isoperimetry. arXiv (2021) doi:10.48550/arxiv.2105.12806.

